# Single and combined effect of salinity, heat, cold, and drought in Arabidopsis at metabolomics and photosynthetic levels

**DOI:** 10.1101/2024.06.23.600276

**Authors:** Elena Secomandi, Marco Armando De Gregorio, Alejandro Castro-Cegrí, Luigi Lucini

**Affiliations:** Department for Sustainable Food Process, Università Cattolica del Sacro Cuore, Via Emilia Parmense 84, 29122 Piacenza, Italy; Department of Sciences, Technologies and Society, University School for Advanced Studies IUSS, Piazza della Vittoria 15, 27100, Pavia, Italy; Department of Plant Physiology, University of Granada, Avenida del Hospicio, 18010 Granada, Spain

**Author notes:** these authors contributed equally to the work (Elena Secomandi, Marco Armando De Gregorio).

**Keywords:** Climate change, multiple abiotic stress, osmolytes, oxidative imbalance, photosynthetic efficiency, secondary metabolism

## Abstract

Ensuring food security is one of the main challenges related to a growing global population under climate change conditions. The increasing soil salinity levels, drought, heatwaves, and late chilling severely threaten crops and often co-occur in field conditions. This work aims to provide deeper insight into the impact of single vs combined abiotic stresses at the growth, biochemical and photosynthetic levels in *Arabidopsis thaliana* L. By studying single and combined stresses, stress interactions and synergic effects have been highlighted. Lower photosynthetic efficiency was recorded from the beginning in all the conditions that included salinity. Consistently, membrane stability and ROS production, combined with a targeted metabolomic quantification of glycine, GABA, proline, and glycine-betaine molecular markers, highlighted the hierarchically stronger impact of salinity and its combinations on plant biochemistry. Untargeted metabolomics coupled with multivariate statistics pointed out distinct metabolic reprogramming triggered by the different stress conditions, either alone or in combination, differentiating the impact of salinity, drought, and their combination with cold and heat. These results contribute to delving into the impact of various stress combinations, hierarchically highlighting the stress-specific effects and pointing out different interactions.

**HIGHLIGHTS:** Combined stresses highlighted synergic and stronger impact on Arabidopsis secondary metabolism, redox imbalance and photosynthetic performance compared to individual stresses. Overall, salinity and its combination were the most impactful.

## INTRODUCTION

Abiotic stresses, such as drought, heat, salinity, and cold, are responsible for a decline in crop yield, both qualitatively and quantitatively (Bashir *et al.,* 2019), and represent worldwide limiting conditions (Cramer *et al.,* 2011). Besides showing differences as a function of stress type, stress level and plant genotype, abiotic stresses can reduce yields by more than 50%, on average (Boyer, 1982; Vogel *et al.,* 2019). Consequently, these stresses are a major threat to food security, becoming even more relevant in the context of an ever-growing human population under climate change. Indeed, climate change exacerbates exposure to abiotic stresses, with extreme temperatures, increased drought, or soil salinity accumulation as the most apparent threats.

The biology of plants’ response to individual abiotic stresses has been extensively studied over the past decades, with several stress-specific modulations (including acclimation) being elucidated at different biochemical and ecophysiology levels. Nonetheless, as reviewed by (Lasky *et al.,* 2023), deeper information about the molecular basis of stress adaptation is still needed. Even more importantly, plants experience a combination of abiotic stresses under realistic field conditions rather than single stresses (Moffat, 2002; Mittler, 2006). Recent literature has pointed out that the response to combined stresses cannot simply be extrapolated from plant response to each individual stress (Rizhsky *et al.,* 2004; Suzuki *et al.,* 2005; Pandey *et al.,* 2015). This suggests that a knowledge gap may exist between the information on plant impact provided by single stresses applied individually compared to multiple stress conditions (Suzuki *et al.,* 2014; Mahalingam, 2015), representing more feasible and realistic field conditions.

Some common biochemical modulations can be observed across abiotic stresses, like the accumulation of osmolytes under drought, salinity, and chilling (Chinnusamy, 2003). On the other hand, plant response to a threat is specific and tailored to environmental stress conditions. A good example of this concept comes from reactive oxygen species (ROS), which are generally associated with most abiotic and biotic stressors but with significant differences in ROS-gene expression patterns observed among different stresses (Mittler, 2002; Mittler *et al.,* 2004). Combined stresses can either be additive or antagonistic to plants (Rillig *et al.,* 2021). For example, conflicting responses are observed when plants open stomata during heat stress to cool leaves by transpiration, thus being more sensitive to co-occurring drought stress. On the contrary, heat stress can increase tolerance to salinity by inhibiting the uptake of Na^+^ ions, promoting their accumulation in roots rather than shoots (Rivero *et al.,* 2014). These few examples make clear that plants’ acclimation to co-occurring abiotic stresses requires a combination of responses to individual stress conditions. Moreover, tailored responses are required to fine-tune molecular processes accounting for the aspects arising from stress combination. Drawing upon the still limited information available in the literature on simultaneously occurring abiotic stresses, Mittler has developed the “stress matrix” where both positive and negative interactions have been proposed and postulated that stress combination should be regarded as a distinct state of abiotic stress (Mittler, 2006).

On these bases, our work aimed at investigating the interaction between co-occurring abiotic stress, using the model plant *Arabidopsis thaliana* L. and applying a combination of biochemical, photosynthetic and metabolomic analyses. The automated, high-throughput phenotyping system monitored plant responses to stresses by evaluating plant growth, leaf shapes, and photosynthetic traits, while the biochemical assays and targeted metabolomic of well-recognized stress markers, together with untargeted metabolomics coupled with multivariate statistics and pathway analysis, aimed at unravelling the additive or antagonistic interactions between drought, heat, chilling, and salinity.

## 2. MATERIAL AND METHODS

### 2.1. Plant material, growing conditions, and stress treatments

Accessions of *Arabidopsis thaliana* L. Columbia-0 (Col-0) were grown from March to April 2023 at the facilities of Università Cattolica del Sacro Cuore (Piacenza, Italy). Seeds were stratified in distilled water and kept for 72 h at 4 °C in dark conditions to synchronize the germination. Pots (6 x 6 x 9.5 cm) were prepared with 130 g of soil:perlite (1:2) mixture and watered one day before sowing up to maximum soil water holding capacity. Then, 5 seeds were sown per pot to be thinned after germination, leaving one seedling per pot. The growth chamber supplied by Ambralight (Ambra Elettronica, Bolzano Vicentino, Italy) was set to 20 ± 2 °C, 8/16 h light/dark photoperiod, and 250 µmol m^-2^ s^-1^ photosynthetic photon flux density (PPFD). The seedlings of *Arabidopsis thaliana* were watered every other day until reaching the 3.7 stage, 38 Days After Sowing (DAS) (Boyes *et al.,* 2001). Plants were then randomly divided into nine groups, each corresponding to a different treatment, with four biological replicates per treatment.

The following treatments were applied: CNTR (Control, unstressed plants), H (Heat), D (Drought), C (Cold), S (Salinity), DxH (Drought x Heat), DxC (Drought x Cold), SxH (Salinity x Heat), SxC (Salinity x Cold). Drought was achieved by blocking irrigation until they reached a relative water content (RWC) of around 70% (9 days). Salinity stress was applied by watering plants with a 100 mM NaCl solution daily until full water holding capacity. Cold stress was induced by keeping the plants at 4 °C for 16 h (dark period). To avoid heat shock, heat stress was applied in two steps: temperature was raised to 26 °C for 14 h and then to 30 °C for 6 h. Both the cold and heat stress were applied the day before the sampling. At the end of the experiments, the leaves collected for metabolomics and biochemical assays were immediately snap frozen in liquid nitrogen to quench metabolisms. Before analysis, samples were ground with pestle and mortar in liquid nitrogen and stored at -80 °C.

### 2.2 RGB imaging and Chlorophyll Fluorescence phenotyping

To investigate the effect of single and combined abiotic stress treatments, plants underwent high-throughput phenotyping measurements. Control and stressed plants were phenotyped for RGB and chlorophyll fluorescence kinetics (ChlF) traits using the PlantScreen^TM^ System (Photon System Instruments, Drásov, Czech Republic). The measurements were conducted starting before the treatment application at 38 and then at 40, 43, 44, 47 DAS (corresponding to T0, T1, T2, T3, T4 respectively). The PlantScreen^TM^ Analyzer software (PSI, Czech Republic) was used to automatically process the raw data (Pixel count and fluorescence intensity) according to Awlia *et al.,* 2016.

RGB images of 5 × 4 plants per tray were captured by using an RGB2 top view camera (GigE PSI RGB, 1.4 Mega-pixels with 1 / 2.3” CMOS SENSOR). Light conditions, plant position and camera settings were fixed throughout the whole experiment. Each round of measurements included an initial 15 min dark adaptation period inside the acclimation chamber.

Chlorophyll fluorescence was acquired using the FluorCam FC-800MF pulse amplitude modulated (PAM) system (PSI, Czech Republic). Three types of light sources were used as part of the ChlF imaging station: (1) PAM short-duration measuring flashes (620 nm), (2) cool-white (6500 K) actinic lights with maximum irradiance 1860 µmol m^−2^ s^−1^ and (3) saturating cool-white light with maximum irradiance 6300 µmol m^−2^ s^−1^. Specifically, to quantify the rate of photosynthesis at diLJerent photon irradiances, an optimization of the Light Curve-Act protocol was applied (Henley, 1993; Rascher *et al.,* 2000) given its suitability to provide information on chlorophyll performances under stress (Brestic and Zivcak, 2013). A 5 s flash of light was applied to measure the minimum fluorescence, followed by a saturation pulse of 800 ms (with an irradiance of 1300 µmol m^−2^ s^−1^) to determine the maximum fluorescence in the dark-adapted state. Next, 60 s intervals of cool-white actinic light were applied at 115, 220, 325, and 430 µmol m−2s−1 corresponding to L1, L2, L3, and L4, respectively. A saturation pulse was applied at the end of the period of actinic light to acquire the maximal fluorescence in the light-adapted state. The ChlF signal measured just before the saturation pulse was taken as the steady-state fluorescence value in the light-adapted state. Fluorescence images were captured by a CCD camera at 16-bit resolution in 1360 x 1024 pixels of CCD chip. Images of various Chl fluorescence ratios obtained by pixel-to-pixel arithmetic operations performed by FluorCam software included maximum PSII quantum yields (F_v_/F_m_), PSII quantum yield of light-adapted plants (F_v_’/F_m_’), coefficient of photochemical quenching (qP) and non-photochemical quenching (NPQ) in steady state.

Finally, the MorphoAnalyser software (version 1.0.9.6) was used to elaborate the acquired images and assess plant growth. The compactness and roundness of leaves were assessed, and the total number of pixels was successively converted to mm^2^ to calculate leaf projected areas.

### 2.2. *Lipid* peroxidation

Lipid peroxidation was determined as malondialdehyde (MDA) content by the TBARS assay (Heath and Packer, 1968; Castro-Cegrí *et al.,* 2023a) with minor modifications. 100 mg of fresh leaves were extracted using 1.5 mL 20% trichloroacetic acid (TCA) (w/v) and 0.3 mL 4% butylated hydroxytoluene (BHT) (w/v). The homogenate was centrifuged twice for 10 min at 10,000 *×* g at 4 °C. 0.25 mL of supernatant was mixed with 0.75 mL of 0.5% thiobarbituric acid (TBA) (w/v); the mixture was incubated during 30 min at 94 °C, then the reaction was stopped in ice for 10 min. The absorbance of the supernatant was then measured at 532 and 600 nm. Results were calculated using a calibration curve and expressed as µg of MDA per kg of fresh weight.

### 2.3. Hydrogen peroxide content

The content of hydrogen peroxide (H_2_O_2_) was determined as in (Alexieva *et al.,* 2001) with minor modifications. 0.4 g of fresh leaves material was homogenized in 1.5 mL of 0.1% (w/v) trichloroacetic acid and centrifuged at 4 °C and 7179 x g for 15 min. The reaction mixture comprised 0.25 mL of supernatant, 0.25 mL of 0.1 M potassium phosphate buffer pH=7 and 1 mL of 1 M KI. Samples were incubated for 1 h in the dark at room temperature, and absorbance was measured at 390 nm. Results were calculated using a calibration curve and expressed as µg of H_2_O_2_ per kg of fresh weight.

### 2.4. Electrolyte leakage

Electrolyte leakage was determined following the method proposed by (Castro-Cegrí *et al.,* 2023b). Each replicate consisted of four leaves of similar size (1 x 3 cm on average), and four replicates were measured per treatment. Leaves were rinsed with 50 mL of deionized water thrice for 3 min. After being incubated for 30 min and shaken at 100 rpm in 30 mL of deionized water, this solution was measured for initial conductivity (Ci) at room temperature using a conductometer. Total conductivity (Ct) was then determined after boiling the flasks for 10 min and cooling at room temperature. The electrolyte leakage was expressed as a percentage of total conductivity: % electrolyte leakage = (Ci*100)/Ct.

### 2.5. Untargeted Metabolomic analysis

An accurate amount (0.200 g) of each sample was extracted using an ultrasonic bath (ArgoLab DU-32; Carpi (MO), Italy) for 15 min at maximum power in 2 mL of 80% methanol (MeOH, purity ≥99.8%, Sigma-Aldrich, St. Louis, MO, USA) solution with 0.1% (v/v) formic acid (purity ≥95%, Sigma-Aldrich, St. Louis, MO, USA). Samples were then centrifuged at 7179 × g for 15 min at 4 °C (Eppendorf 5430R, Hamburg, Germany), and 1 mL of the resulting supernatant was transferred in a vial using a 0.22 µm regenerate cellulose filter. Four independent replicates were analyzed for each treatment condition, with two technical replicates per sample. Quality Controls (QC) were prepared by mixing 20 µm of each extract and were randomly injected throughout the chromatographic sequence to avoid analytical bias.

The phytochemical profile was evaluated by ultra-high-pressure liquid chromatography equipped with a binary pump and a Dual Electrospray Jetstream ionization source, coupled to quadruple time of flight mass spectrometry (1290 UHPLC / 6550 iFunnel QTOF-MS from Agilent Technologies, Santa Clara, CA, USA) as previously reported by Salehi *et al.,* 2023. A volume of 6 μL was injected and reverse phase chromatography was applied for separation using a water-acetonitrile gradient elution from 6 to 94% of acetonitrile in 33 min, a flow rate of 0.2 mL/min and an Agilent Zorbax Eclipse Plus C18 analytical column (15 cm x 2.1 mm, 1.7 µm particle size). The mass spectrometer acquired data in SCAN mode in the 100–1200 m/z range, with a nominal resolution at 30,000 full-width half maximum (FWHM). The QTOF mass analyzer operated in positive mode (ESI+) for both MS and MS/MS acquisition with nitrogen as both sheath gas (12 L/min and 315 °C) and drying gas (14 L/min and 250 °C). The nebulizer pressure was 45 psi, the nozzle voltage was 350 V, and the capillary voltage was 4.0 kV for MS acquisition. Blank samples were injected at the beginning and the end of each randomized sequence run. Moreover, QC samples were analyzed within each sequence every 9 samples in data-dependent MS/MS mode (8 precursors per cycle, 1 Hz, 50–1200 m/z, positive polarity, active exclusion after 2 spectra), at 10, 20, and 40 eV collision energies.

Raw data were annotated by Profinder B10.0 (Agilent Technologies) applying the “find-by-formula” algorithm based on monoisotopic mass (5-ppm tolerance for mass accuracy), isotope spacing and ratio. Compound annotation was carried out against the PlantCyc database 9.6 (Hawkins *et al.,* 2021) following mass and retention time alignment of deconvoluted features. Level 2 of identification (i.e., putatively annotated compounds, COSMOS standards in metabolomics) was achieved (Salek *et al.,* 2013). Data filtering was finally applied, and compounds that were not detected in at least 75% of the replications within at least one group were discarded.

### 2.6 Targeted analysis for osmolyte quantification

A targeted approach was used to quantify the osmolyte stress markers proline, glycine, glycine-betaine, and γ-aminobutyric acid (GABA). To this aim, a Vanquish ultra-high-pressure liquid chromatography coupled to a Q-Exactive HF Hybrid Quadrupole-Orbitrap mass spectrometer through a HESI-II probe (Thermo Scientific, Waltham, MA, USA) was used. The chromatographic and MS conditions were based on Khan *et al.,* 2017. Briefly, a volume of 6 μL of the same extract used for untargeted metabolomics was injected. The separation was made by a water-acetonitrile gradient elution of 50% in 12 min with a flow rate of 0.2 mL/min at 50 °C using an Acquarity PREMIER Peptide CSH C18 analytical column (130A 1.7 µm, 2.1 x 150 mm, 1/pk). The mass spectrometer acquired data in FULL SCAN mode in the 50–250 m/z range, with a nominal resolution at 70,000 full of at half maximum (FWHM) and in positive polarity. The Orbitrap mass analyzer operated in positive mode (ESI+) for both MS and MS/MS acquisition with nitrogen as both sheath gas (40 L/min) and auxiliary gas (20 L/min and 50 °C). The spray voltage was 3.5 kV with a capillary temperature of 250 °C and an S-lens RF level of 50. XCalibur 4.1.31.9 (Thermo Fisher Scientific Inc.) software was used for data acquisition and processing. Absolute quantification was achieved against calibration curves built with pure reference standards for proline, glycine, betaine, and GABA (all from Sigma-Aldrich; purity >98%). Monoisotopic accurate mass, MS/MS spectral fragmentation, and retention time were used for identification (Supplementary Table S1). All quantitative results were expressed as μmol g^−1^ FW apart from betaine data that were expressed as nmol g^−1^ FW.

### 2.7 Statistical analysis

One-way ANOVA with Duncan post-hoc test (*P*<0.05) was carried out for morpho-physiological and photosynthetic data, biochemical assays and osmolytes quantification using SPSS 28 (IBM, Armonk, NY, USA). Linear correlation was performed on the same software.

Metabolomics data were elaborated with Mass Profiler Professional B15.1 software tool (Agilent Technologies, Santa Clara, CA, USA) (Benjamin *et al.,* 2019). Compounds abundance was log2-transformed, normalized at the 75^th^ percentile and baselined against the median of all samples. According to their metabolic profile, the similarities and/or differences among samples were reported through unsupervised hierarchical cluster analysis (HCA - Euclidean distance, Ward’s linkage) based on fold change (FC) heat map. One-way analysis of variance (ANOVA) was performed to detect the statistically significant compounds among treatments, setting a significance level of *P*<0.05 (Tukey’s post hoc test; Bonferroni multiple testing correction). Datasets were separately imported in SIMCA 17 (Umetrics, Malmo, Sweden) to perform supervised orthogonal projection to latent structures discriminant analysis (OPLS-DA) multivariate modelling. The obtained model was successively cross-validated (CV-ANOVA; *P*<0.05), inspected for outliers (Hotelling’s T2), and the model’s goodness parameters (goodness-of-fit R^2^Y and goodness-of-prediction Q^2^Y) were checked. Model overfitting was excluded by permutation testing (n=100). Finally, the Variable Importance in Projection (VIP) analysis was used to select the metabolites having the highest discriminant potential score (VIP>1.2). Statistically significant (*P*<0.05 in at least one of the treatments) VIP compounds were selected for biochemical interpretations in Pathway Tools Omic Dashboard (version 27.0) (Plant Metabolic Network, http://www.plantcyc.org, accessed on November 24^th^, 2023) (Paley and Karp, 2024).

## 3. RESULTS

### 3.1 Growth *and morphological traits*

Phenotyping allowed the non-destructive monitoring of plant growth throughout the entire experimental period. Data regarding the plant area showed no significant differences in the first part of the experiment, while from T3, corresponding to the 6^th^ day of salinity, stress treatment reduced plant growth and all the salt-stressed plants were significantly different from the non-salinity stressed ones (Fig. 1A; Supplementary Table S2). However, heat and cold stress applications did not impact the plants’ area, likely because of the limited duration of the stress. Similarly, drought stress didn’t affect the digital area of plants.

**Fig. 1.**
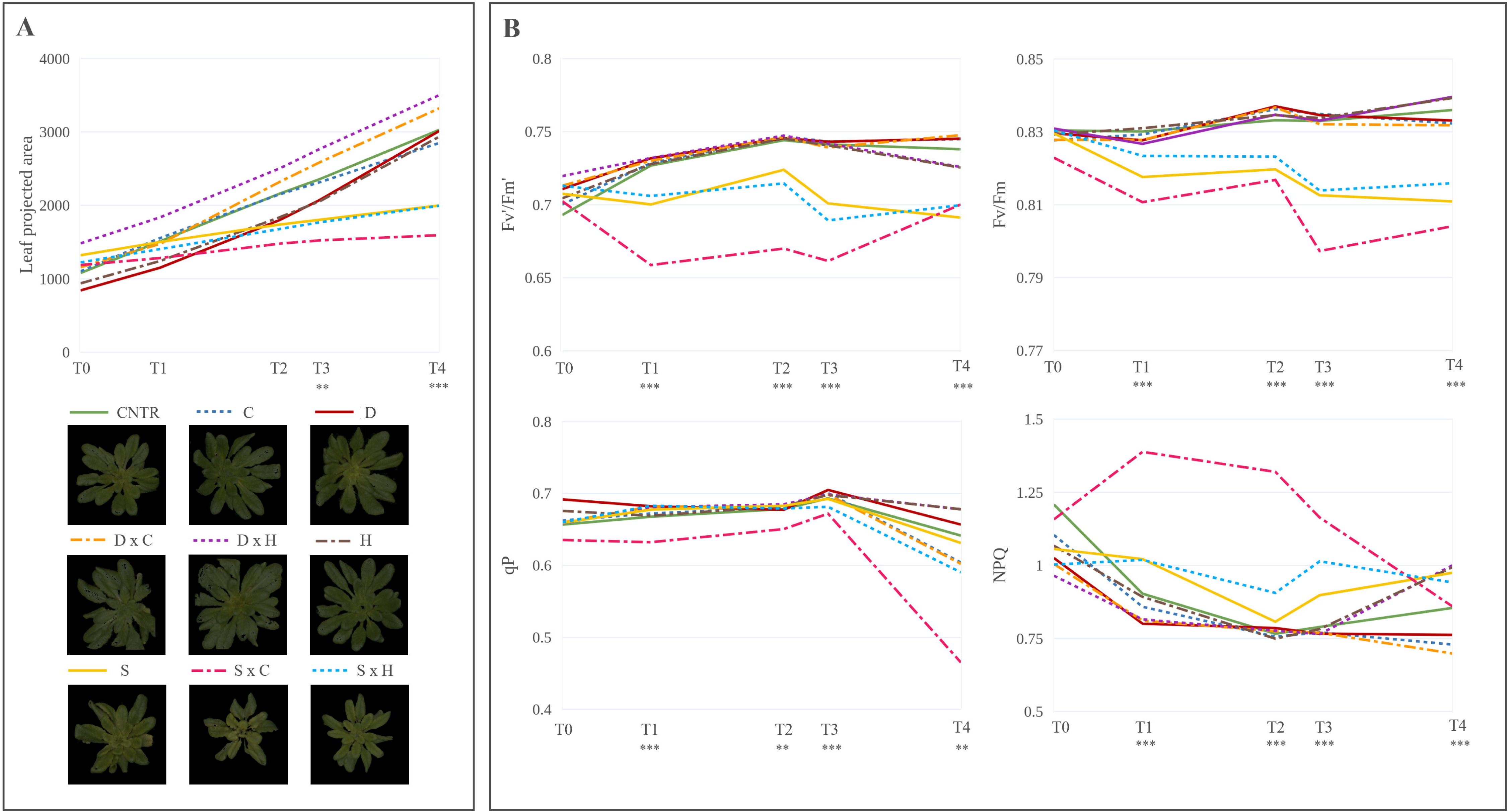
**(A)** Leaf Projected Area of *Arabidopsis thaliana* under control, single and combined abiotic stresses and RGB images of plants at T4 **(B)** Maximum quantum yield of PSII in dark-adapted (F_v_/F_m_) and light adapted (F_v_’/F_m_’) leaves, non-photochemical quenching (NPQ) and coefficient of photochemical quenching in steady state (qP) of *Arabidopsis* plants under control, single and combined stress conditions. All data were acquired with the PlantScreen System; T0, T1, T2, T3, T4 correspond to 38, 40, 43, 44, 47 DAS. Significant differences between control and stress treatments are indicated with ** and *** for *P*<0.01 and 0.001, respectively.

Other morphological parameters, including compactness and roundness, have been further addressed. In the late phase, significant differences for roundness could be observed in SxH-treated plants while compactness never differed in the T0-T4 measurements (Supplementary Table S2).

### 3.2 Photosynthetic *performance*

To determine the physiological status of *Arabidopsis* plants under single and combined stress conditions, an automated chlorophyll fluorescence imaging was set based on the light curve protocol. The basic ChlF parameters were derived from the measured fluorescence transient states (e.g., F_0_, F_0_’, F_m_, F_m_’, F_t_, and F_v_) and then used to calculate the quenching coefficients (e.g., qP, NPQ) and other plant photosynthetic performance parameters (e.g., F_v_/F_m_, F_v_’/F_m_’). The maximum quantum yield of PSII photochemistry in the dark-adapted (F_v_/F_m_) and the light-adapted (F_v_’/F_m_’) states, the coefficient of photochemical quenching that estimates the fraction of open PSII reaction centers (qP), the steady-state non-photosynthetic quenching measuring heat dissipation (NPQ) for days 0, 2, 5, 6, and 9 of the stress-application periods are showed in Fig. 1B. The highest actinic photon irradiance (L4) was chosen to assess the photosynthetic activity variation as it provided the most discriminative power for evaluating abiotic stresses impact on ChlF parameters.

The analysis of NPQ and qP parameters allowed to highlight interesting and diverse responses in the photosynthetic activity amid treatments. Upon exposure to salt stress, increased NPQ values were observed at varying degrees in salt-treated plants starting from T1. At T4, after the temperature variation inductions, a strong increase could be recorded in heat-treated plants (Fig. 1B; Supplementary Table S3). Regarding qP, a decrease was observed in the salt-induced over time and in the cold-treated plants at T4. Overall, considering the longer salt stress applications, this resulted in a rapid and substantial increase in non-photochemical processes (i.e., the dissipation of heat in the PSII antennae), which correlates with reduced PSII quantum efficiency and photochemical quenching under stress (Fig. 1B; Supplementary Table S3). However, temperature induction resulted in a variation of NPQ and qP, respectively, for heat and cold stress, without impacting the overall F_v_/F_m_ values, likely due to the short time application. On the contrary, a progressive decline of the maximum quantum yield (F_v_/F_m_) was recorded in plants under salinity starting from T1 (Fig. 1B; Supplementary Table S3). No significant change in the photosynthetic efficiency could be observed in drought-stressed plants during the 9 days of stress, suggesting no damage to PSII throughout the stress treatment. Comparable results were recorded for F_v_’/F_m_’, which did not change between control and stressed plants, except for salt-treated samples.

#### 3.2.2 Effect of combined stresses on oxidative stress markerss

Abiotic stresses can increase reactive oxygen species, such as hydrogen peroxide (H_2_O_2_). As shown in Table 1, salinity and its combinations significantly increased the accumulation of H_2_O_2_ compared to the control, with the combination of salinity and cold stress being the treatment triggering the highest accumulation of this compound. Comparing the results obtained from MDA content as an indicator of lipid peroxidation, all the stresses induced MDA accumulation, compared to control (Table 1). Furthermore, the salinity stress and its combinations showed the highest induction of this parameter than the other stresses, particularly for SxC, where an accumulation of 388% was recorded compared to the control (Table 1). The interaction between MDA and lipid peroxidation was evaluated through linear correlation analysis, showing a Pearson’s correlation factor of 0.915 (*P*<0.001) between the two stress markers. When analysing membrane stability, salinity was consistently the stress that affected leaves the most, damaging this tissue and causing up to a 360% increment in electrolyte leakage compared to the control (Table 1).

**Table 1.**
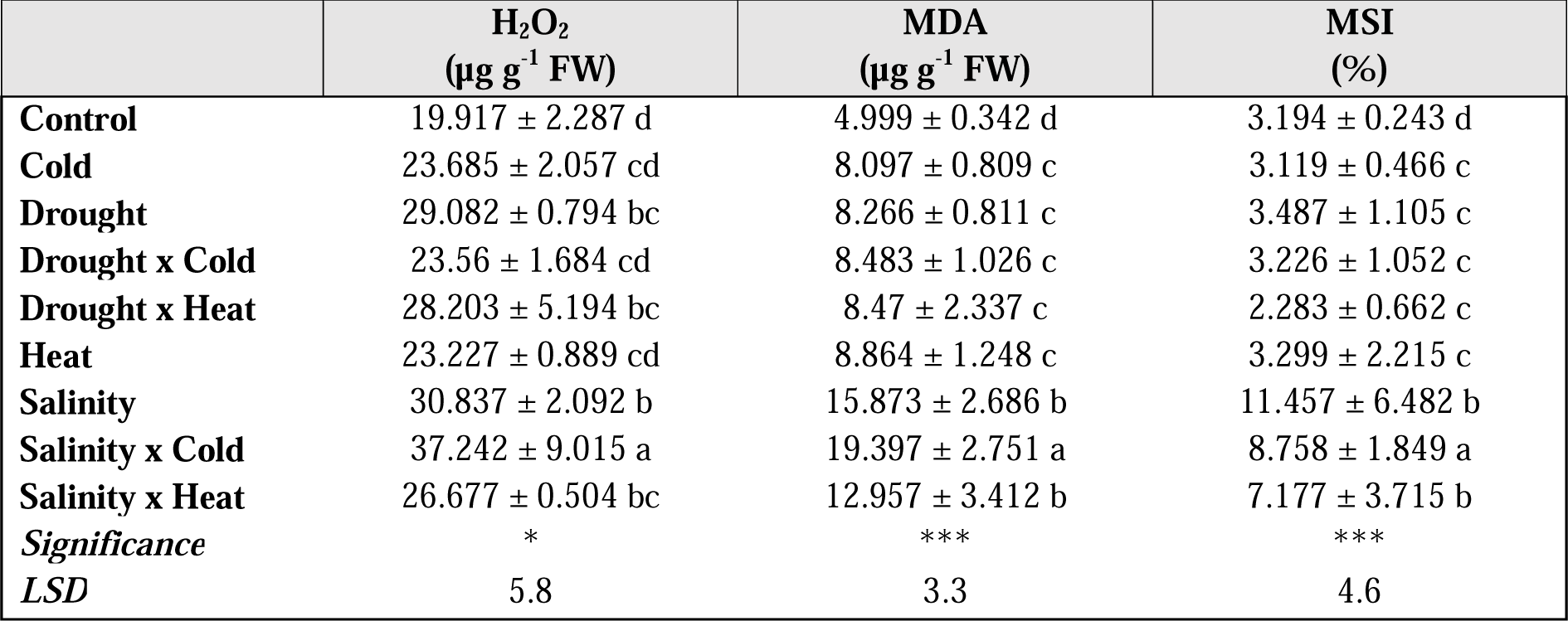
Changes in peroxide oxide (H_2_O_2_, µg g^-1^ FW), malondialdehyde (MDA, µg g^-1^ FW) and Membrane Stability Index (MSI, %) in control plants and plants under single (salinity, drought, heat and cold) stresses and their combination. Data represent the mean ± standard deviation of 4 plants. Different uppercase letters indicate differences between treatments after one-way ANOVA with Duncan’s post hoc test, while asterisks indicate significant differences (* *P*<0.05; *** *P*<0.001).

### 3.3 Metabolomics *profile of leaves*

An untargeted metabolomics approach was used to investigate the effect of different stresses and their combinations on the metabolomic profile of *Arabidopsis* leaves. This approach allowed us to annotate more than 2000 putative compounds in leaves. The list of compounds, together with individual raw data abundance, retention time and composite mass spectra are reported in Supplementary Table S4, whereas the raw data are published in the repository MetaboLights (Yurekten *et al.,* 2024) under study ID MTBLS9669.

First, unsupervised hierarchical cluster analysis was performed to investigate patterns across the conditions considered naively and to provide a hierarchical picture of the factors under study (Fig. 2). As expected, the heat map based on fold-change highlighted a distinct metabolomic profile depending on the stress applied and provided a hierarchical overview of the different conditions. Two main clusters are visible, the first including control and single stresses, the second including salinity single stress and all the combined stresses, with drought and salinity-related stresses clustering together.

**Fig. 2.**
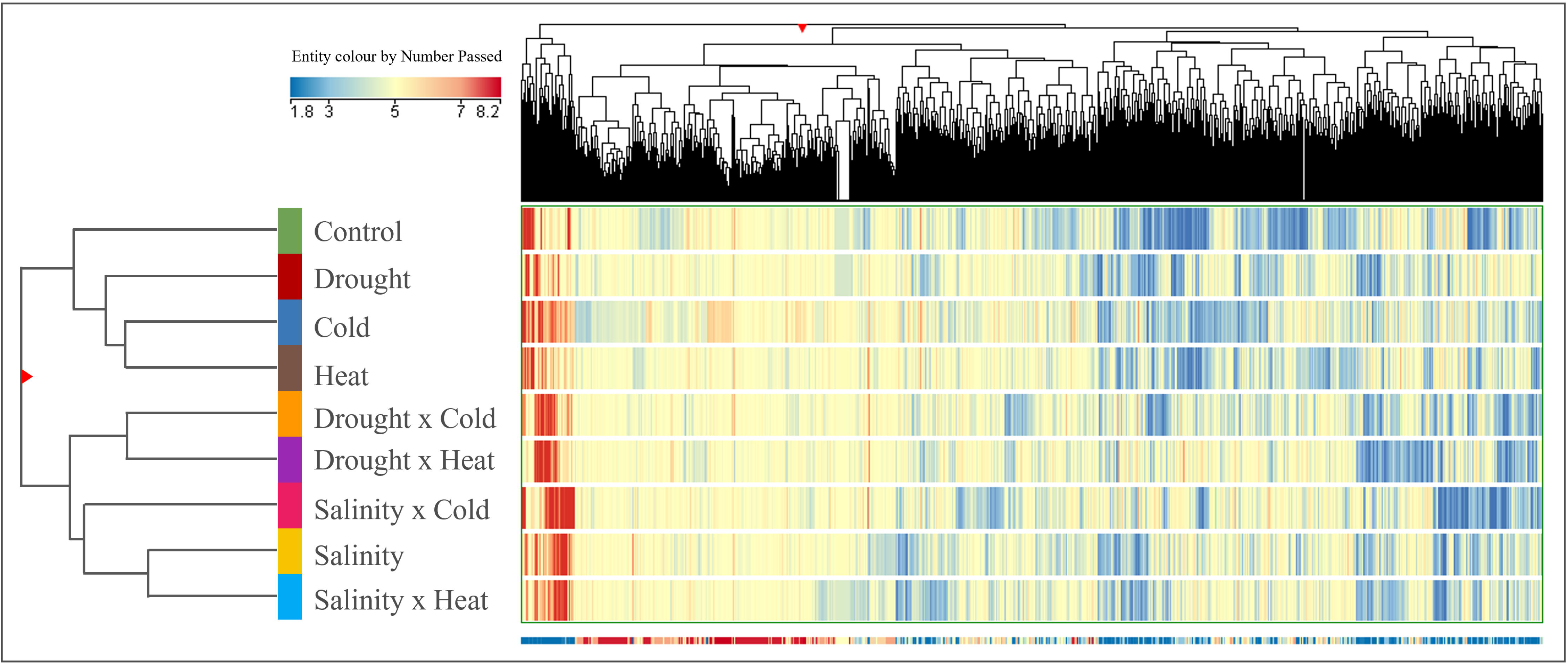
Unsupervised HCA of the metabolomic profile obtained through UHPLC-ESI/QTOF-MS analysis of Arabidopsis grown under control, single stress conditions (e.g., cold, drought, heat, salinity) and their combination. Hierarchical clusters (linkage rule: Ward; distance matrix: Euclidean) were based on the fold-change-based heat map of compounds’ normalized intensities.

These results were further investigated by the supervised OPLS-DA based on all treatments (Supplementary Fig. S1), corroborating the HCA unsupervised clustering outcomes. The model presented adequate scores regarding the goodness-of-fit (R^2^Y) = 0.957 and prediction ability (Q^2^Y) = 0.735. Based on the OPLS-DA discriminating model, a Variable Important in the Projection (VIP) analysis was carried out to identify the compounds having the highest discriminant power (VIP score > 1.2, *P*<0.05). Overall, 255 compounds were found to be discriminating by the OPLS-DA model. Among these, the most abundant classes included polyphenols (34), terpenes (14), nucleosides (14), alkaloids (11) and glucosinolates (6). The full list of statistically significant VIP markers is available in Supplementary Table S5.

Volcano analysis (FC>2, *P*<0.05) highlighted 566 compounds being statistically different from the control in at least one treatment (Supplementary Table S6). Specifically, for individual stresses, the number of differential metabolites was 132 for C, 71 for H, 166 for D and 259 for S. Differential compounds that differed under combined stresses were 204 for DxC, 313 for DxH, 244 for SxC and 274 for SxH. The selected compounds were then plotted on the PlantCyc pathway tool to understand better the metabolic impact driven by the different stresses applied compared to the control (Supplementary Fig. S2). According to the pathway tool, drought and salinity treatments and their combinations induced a stronger modulation of secondary metabolites, hormones, and amino acid biosynthesis. Among secondary metabolites, the classes showing the stronger modulation were N-containing compounds (including glucomalcommin; S-magnoflorine; caffeoylserotonin), S-containing compounds (such as methiin and gamma-L-glutamyl-(S)-methyl-L-cysteine) and Terpene-related compounds (such as acetoacetyl-CoA; 10-deoxysarpagine; phytyl diphosphate). Concerning amino acid biosynthesis, contrasting trends could be observed: while Arg, Glu, Ile, Lys, Pro and Thr were accumulated, Phe, Trp, and Tyr decreased in all conditions. Among hormones, melatonin, and Gibberellin A38 decreased in all considered conditions. In DxH and SxC combined stresses, 2-*cis*,4-*trans*-xanthoxine was positively accumulated, while in DxH, SxH and SxC interaction, the *trans*-zeatin riboside triphosphate increased. These results suggest a different stress response mainly due to the stress type.

To delve into the compounds driving this separation, two OPLS-DA models were developed based on salinity and drought stresses and their combination (Fig. 3A and 3B). In both cases, the models presented adequate scores (R^2^Y = 0.991 and Q^2^Y = 0.831 for S stress and its combinations and R^2^Y = 0.996 and Q^2^Y = 0.856 for D stress and its combinations). The VIP analysis (VIP score threshold = 1.2, *P*<0.05) was then carried out, and the full list of statistically significant VIP markers under drought and salinity stresses and their combinations is provided in Supplementary Tables S7 and S8, respectively). Overall, 406 compounds were found as discriminants in the OPLS-DA model based on drought stress and its combinations. Among these, polyphenols (34), terpenes (25), alkaloids (18), indole derivates (15), fatty acids (14), glucosinolates (13) and phenylpropanoids (9) were included.

**Fig. 3.**
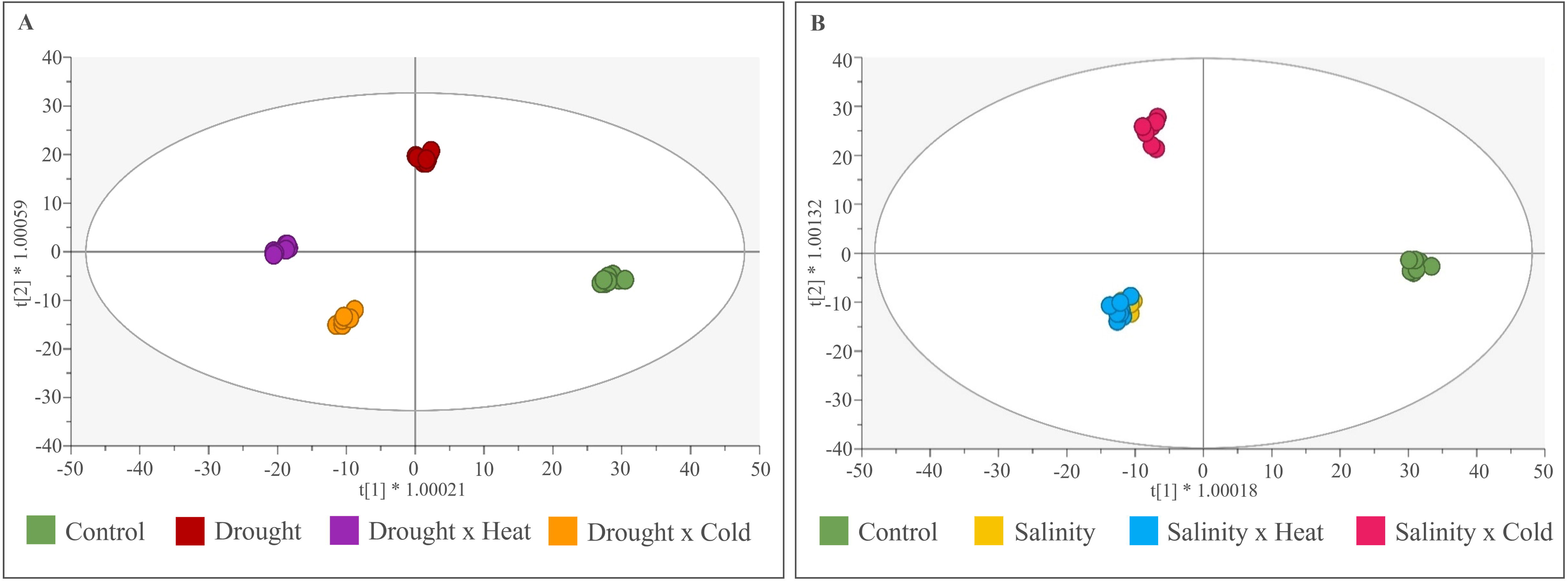
Supervised OPLS-DA based on the metabolomic profile of Arabidopsis leaves under (A) drought condition and its combination (R2Y = 0.996; Q2Y = 0.856) and (B) salt condition and its combination (R2Y = 0.991; Q2Y = 0.831). The VIP markers (VIP-score > 1.2) extrapolated by the models are listed in Supplementary Table S7 and S8, respectively.

Concerning the OPLS-DA model based on salinity stress and its combinations, 405 discriminant compounds were identified. Among these, the most represented classes were polyphenols (32), amino acid derivates (29), terpenes (28), alkaloids (19), indole derivates (14) and fatty acids (11).

To identify the metabolite in common between drought and salinity single stresses and their respective combination, Venn Diagrams were elaborated on compounds with FC>2 and *P*<0.05 (Fig 4 and Supplementary Fig. S3). For drought and its combinations, 83 common compounds were identified, 45 of which had a VIP-score>1.2 (Supplementary Fig. S4A); when considering all the five stress conditions, among the 23 compounds in common, 9 had a VIP score>1.2 (Fig 4A). Under salt applications, 132 compounds were common across S, SxC and SxH; among them, 71 had a VIP-score>1.2. When considering also heat and cold single stresses, only 7 compounds out of the 19 in common to all the five conditions had a VIP-score>1.2 (Fig. 4B).

**Fig. 4.**
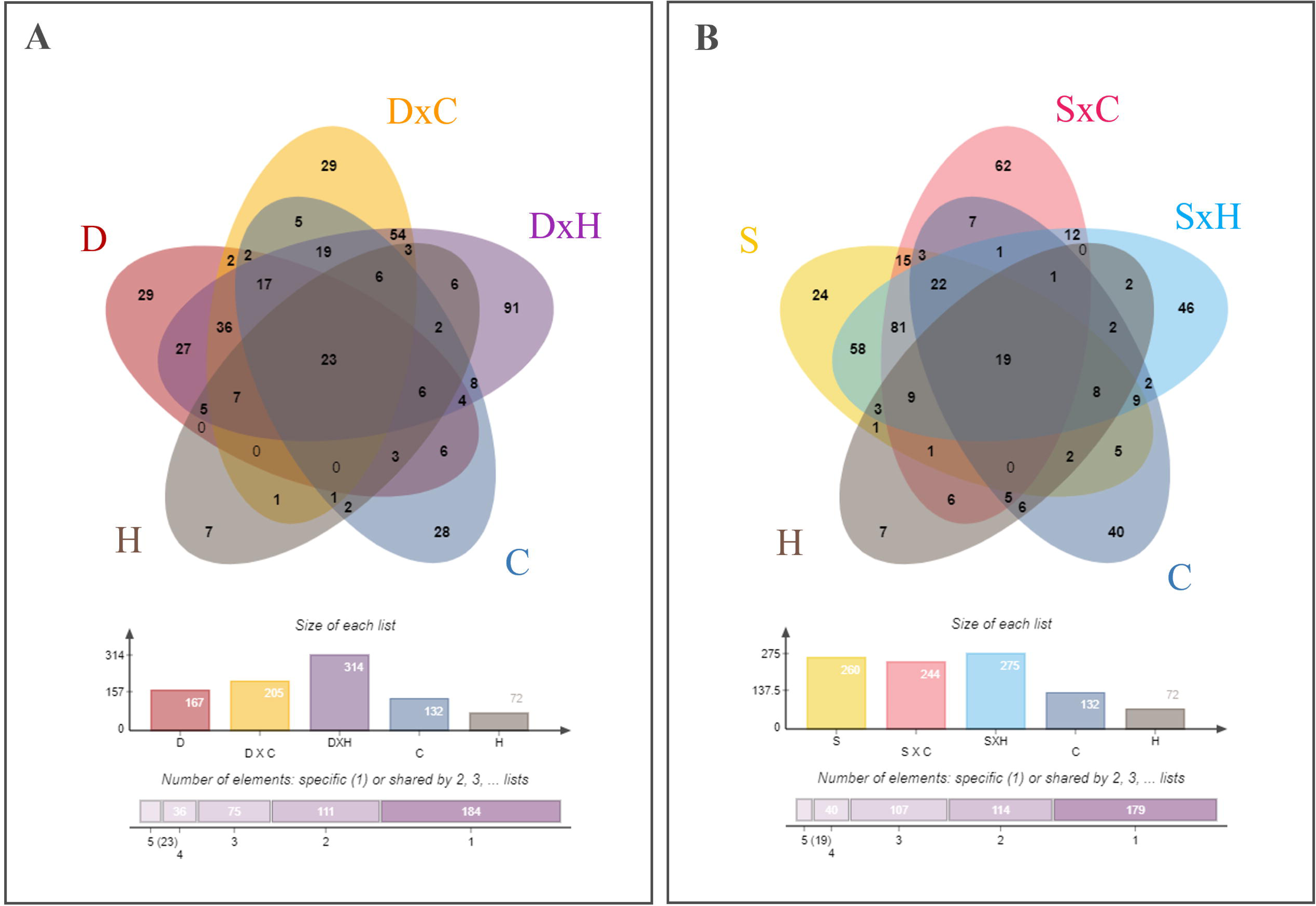
Venn diagrams comparing the differential metabolites resulting from the Volcano analysis (P<0.05; FC>2) in (A) plants under drought condition and (B) under salinity stress.

### 3.4 Impact *of multiple stresses on osmolytes*

A targeted approach identified and quantified key osmolytes involved in stress response, including GABA, proline, betaine, and glycine. Data regarding RT, MS1 and MS/MS spectra are reported in Supplementary Table S1. In our experiment, the applied abiotic stresses elicited proline and betaine accumulation (Table 2). Salinity and its combinations significantly increased the accumulation of proline compared to the control, with SxH being the treatment with a stronger effect (14.5-fold increase). On the opposite, betaine levels were lower in stressed samples. Specifically, compared to the control, all single stressed plants apart from salinity-treated samples and all drought combinations showed a reduction in this compound.

**Table 2.**
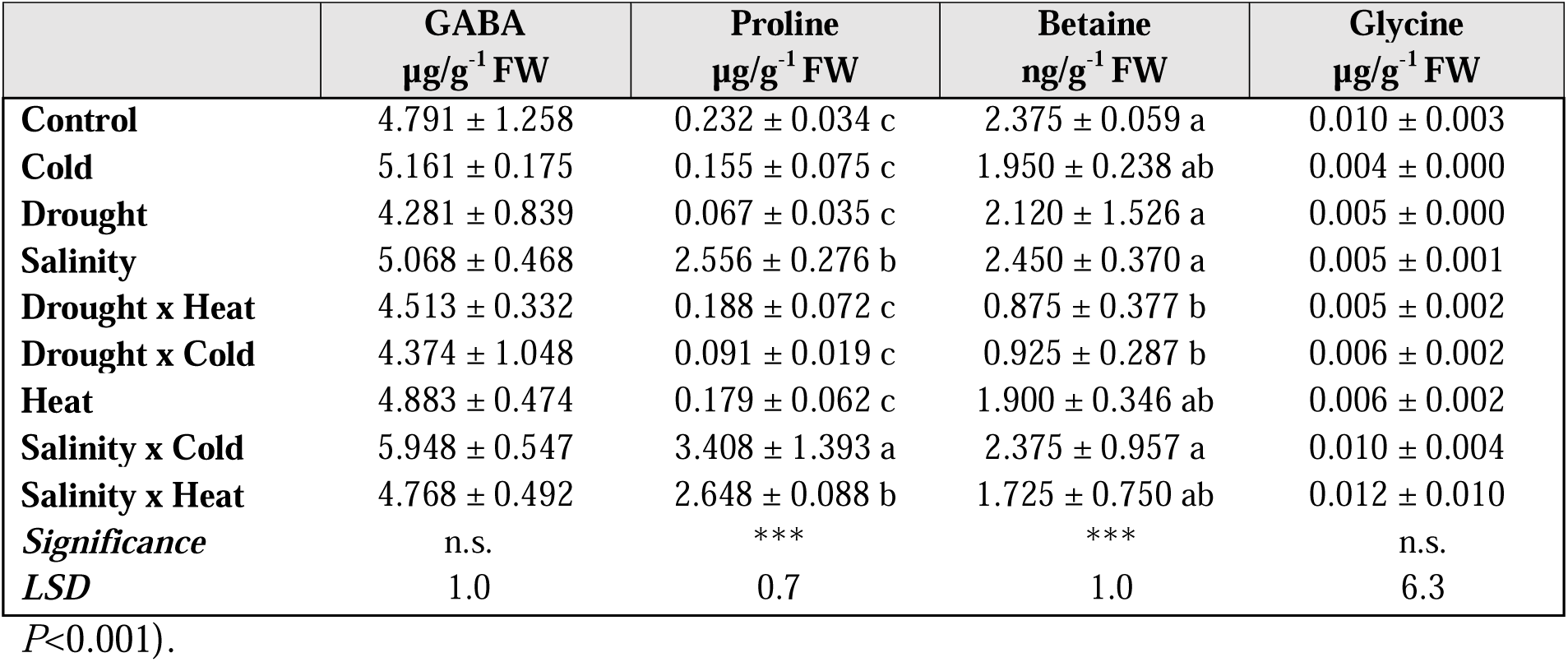
Changes in γ-aminobutyric acid (GABA, µg g^-1^ FW), proline (µg g^-1^ FW), betaine (ng g^-1^FW) and glycine (µg g^-1^ FW) in control plants, and plants under single stresses (salinity, drought, heat and cold) and their combination. Data represent the mean ± standard deviation of 4 plants. Different uppercase letters indicate differences between treatments after one-way ANOVA with Duncan’s post hoc test, while asterisks indicate significant differences (n.s., non-significant; *** *P*<0.001).

## 4. DISCUSSION

By impacting plants, the rising number and severity of extreme climate change-related events heavily threaten food production worldwide (Mbow *et al.,* 2019). To face these changing environmental conditions, plants respond by adapting their physiology through widespread changes in cellular processes (Zhang *et al.,* 2022). The coping mechanisms for stress include a shift in photosynthetic performance, accumulation of osmoprotectants and antioxidants, and phytohormone modulation (Gupta and Huang, 2014; Ondrasek *et al.,* 2022). These processes finally result in a change in plant growth rate. However, this extensive metabolism shift represents an intricate network of responses that overlap and crosstalk, thus being sometimes difficult to understand.

The effective evaluation of plant growth and performance has been assessed using phenomics as a rapid and non-invasive technique representing physiological performance to unfold the plant’s response under these fast-changing conditions. Indeed, in plant biology, chlorophyll fluorescence is widely recognized to investigate Photosystem II (PSII) efficiency and to describe responses to environmental changes (Schurr *et al.,* 2006), including stresses (Chaerle and Van Der Straeten, 2000).

In our experiments, RGB images were poorly predictive since they revealed that abiotic stresses didn’t affect rosette morphology in *Arabidopsis*. Different results could likely be observed in long-term experiments. Regarding growth rate, salt-stressed plants significantly reduced leaf area from the late phase of the treatments (T3-T4). On the contrary, temperature stresses were applied as a single time point, thus without influencing biomass. Surprisingly, no impact on plant growth could be observed under drought conditions, probably due to the phenological phase at induction, the duration, and the severity of stress conditions. Indeed, in other experiments on *Arabidopsis*, plant growth was affected when longer drought periods were applied, suggesting a duration-dependent dose before the impact on biomass turns visible. Alongside this, the phenological phase when stress is applied can be pivotal in defining the overall impact, with a stronger effect on seedlings or young plant (Chaerle and Van Der Straeten, 2000; Harb *et al.,* 2010; Vile *et al.,* 2012). Overall, these results on plant growth suggest that this parameter may not be an ideal indicator of stress, even more in multiple stress conditions.

According to previous research (Shahid *et al.,* 2020; Giannelli *et al.,* 2023), a reduced growth rate can correlate to lower photosynthetic performance. In fact, since the T1 corresponded to a 2-day salinity treatment, salt application significantly affected the PSII efficiency, with a reduction in all photosynthetic parameters (Fig. 1; Supplementary Table S2). Nevertheless, the recorded values showed interesting results related to photosynthetic performance for most of the treatments. Indeed, the different stresses distinguishingly impacted the non-photochemical processes, represented by the NPQ parameter, and the photochemical efficiency, represented by the F_v_’/F_m_’ and qP traits. Specifically, while salinity affected all these parameters, possibly due to its duration, cold and heat stresses had a distinct effect on the photosynthetic apparatus, turning into an increase in the NPQ mechanism of heat dissipation in heat-stressed plants and a decrease in the number of the PSII open centres in cold-treated samples.

These parameters quantifying photochemical and non-photochemical processes display more dynamic responses to stresses, eventually being followed by a reduction in the maximal quantum efficiency of PSII in dark-adapted state (F_v_/F_m_; Mishra *et al.,* 2012). Although the maximum quantum yield is commonly used for evaluating the performance of stressed plants (Jansen *et al.,* 2009; Bresson *et al.,* 2015), F_v_/F_m_ is an extremely stable parameter that often cannot be used to monitor early stress in agreement with previous reports (Baker and Rosenqvist, 2004; Kalaji *et al.,* 2017). In our experiment F_v_/F_m_ appears to be a robust parameter (Fig. 1; Supplementary Table S3), being affected only under prolonged stress conditions (salinity) and not reflecting short stress responses (e.g., cold and heat). Interestingly, despite the 9-day drought application, any impact on the photosynthetic apparatus could be recorded, suggesting that an overall mild stress level was reached on plants. As already reported (Suresh *et al.,* 2012), the suitability of Chl *a* fluorescence parameters for assessing drought depends on the severity and duration of the stress. Despite causing a decrease in the photosynthetic rate, mainly due to stomatal closure, mild-to-moderate drought stress has no direct effect on the individual metabolic reactions’ capacity (Brestic *et al.,* 1995; Cornic and Massacci, 1996; Flexas and Medrano, 2002). Notwithstanding, drought stress may worsen the effect of co-occurring stresses (e.g., qP in DxC and NPQ in DxH samples, Supplementary Table S3).

Phenotyping investigations revealed the lack of single unique markers suitable for investigating stress interactions. Consequently, untargeted metabolomics was then performed to delve into the effect of single and combined stress on the overall metabolism beyond hypothesis-driven specific stress markers. The unsupervised hierarchical analyses revealed the distinctive effect of single and combined stress on plant metabolism, confirming the necessity to unravel the intricate responses underlying the different stress conditions. This was also confirmed by the Venn analyses on differential metabolites under salinity or drought stress, compared to their combination with heat and cold. The results (Fig. 4) pointed out the lack of shared markers across all the stress conditions and rather pointed out a stress-tailored response to each condition. On the contrary, stress combinations shared more similar biochemical responses with their respective single stress, while keeping their own singularity in the plant metabolomic signature, thus acting as a new stressor (Mittler, 2006; Zandalinas *et al*., 2021).

The analysis of the significant compounds (*P*<0.05) with an FC>2 highlighted the main categories of metabolites affected by the treatments compared to the control. These include an accumulation of glucosinolates, alkaloids and polyphenols, all involved in oxidative stress mitigation. Specifically, glucomalcommin is linked to the production of aliphatic glucosinolates in water stress response (Zhang *et al.,* 2021), and in general, glucosinolates have been correlated to the aquaporins modulation (Martínez-Ballesta *et al.,* 2015). S-magnoflorine is an isoquinoline alkaloid with an anti-oxidative effect against the oxidation of lipoproteins (Hung *et al.,* 2007; Okon *et al.,* 2020), while caffeoylserotonine is a precursor of melatonin (Byeon *et al.,* 2014). Melatonin, considered a novel phytohormone, is involved in plant growth and tolerance to abiotic stress by directly scavenging free radicals, enhancing ROS detoxification, and regulating the enzymatic and non-enzymatic antioxidant systems (Zhang *et al.,* 2020).

Among S-contain compounds, the metabolites related to the GSH detoxification system and derivates of glutathione, which plays an important role in controlling cellular redox balance and detoxifying side products of plant metabolism, were represented. Terpenes are the last class of compounds mostly affected by stress-induced changes. In fact, terpenes have a broad set of biological functions (Dudareva *et al.,* 2006) and are involved in various stress-induced responses (O’Connor and Maresh, 2006). In particular, acetoacetyl CoA is an upstream metabolite in the mevalonic acid pathway (Ahumada *et al.,* 2008). The down accumulation of phytyl diphosphate, involved in the phythol pathway, can be related to the impact of abiotic stresses on photosynthetic activity, being related to chloroplast’s structural changes (Gutbrod *et al.,* 2019). While the thylakoids membrane decreases, starch granules increase (Gaude *et al.,* 2007), with alterations in the lipid ultrastructural composition (Tevini and Steinmüller, 1985), thus resulting in the release of large amounts of phytol due to chlorophyll degradation (Lippold *et al.,* 2012). Interestingly, Ischebeck *et al.,* 2006 demonstrated that the free phytol produced by the hydrolysis of chlorophyll can, in stress response, be reincorporated into chlorophyll or be used for the synthesis of tocopherol and phytyl esters of fatty acids.

In general, the metabolomic analysis highlighted a reduction in the accumulation of photosynthetic pigment precursors under stress. Moreover, in OPLS-DA analysis of salinity, drought, and their combinations, protochlorophyll and 3,8-divinyl protochlorophyllide-a + chlorophyll-a were highlighted in the two models, respectively.

In general, the maintenance of membrane stability has a key role in plant response to the environment and therefore cell wall components and the proteo-lipid fractions are subjected to a thigh control. Indeed, membrane stability depends on its lipid composition, which controls membrane fluidity (Wang *et al.,* 2020; Rawat *et al.,* 2021). Abiotic stresses can trigger lipid peroxidation due to increased reactive oxygen species. Accordingly, all the abiotic stresses evaluated in this experiment resulted in a higher MDA content. Interestingly, salinity stress had the highest impact, probably due to the significant increment observed in hydrogen peroxide content compared to the other stresses, which correlated with MDA. Overall, the reported increase in reactive oxygen species and lipid peroxidation led to an important loss of membrane integrity in the plants exposed to stress by salinity.

Frequently, osmotic stress in plants under abiotic stress conditions is caused by ion imbalance and water deficiency. Some of the most studied effects on biophysical changes are reduction in cell turgor pressure, shrinkage of the plasma membrane, and physical alteration of the cell wall (Park *et al.,* 2016). In response to stresses, osmolytes such as proline, glycine-betaine and sugars are produced and accumulated as non-toxic molecules (Chen and Murata, 2002). Osmolytes are mainly involved in regulating osmotic pressure but may also influence ABA levels and gene expression; although the response to osmotic stress is fundamental, it impacts plant growth (Giannelli *et al.,* 2024).

Proline is one of the most important osmolytes and plays a role in stabilizing sub-cellular structures, detoxifying ROS, buffering cellular redox potential, and stabilizing protein and protein complexes under stress conditions (Muchate *et al.,* 2016; Sánchez *et al.,* 2023). Proline is synthesized mainly from glutamate in plants, but intracellular proline levels are determined by biosynthesis, catabolism and transport between cells and different cellular compartments (Szabados and Savouré, 2010). Proline biosynthesis is activated by the activity of 2 *P5CS* genes; in *Arabidopsis*, the isoform *P5CS*1 is induced by osmotic and salt stresses (Yoshiba *et al.,* 1995). Székely *et al.,* 2008 through an experiment conducted with GFP, demonstrated that P5CS1-p under stress was located from cytosol to chloroplast. During stress conditions, the Calvin cycle is inhibited, and the oxidation of NADPH is reduced; therefore, the flow of electrons in the electron transport chain is interrupted and ROS accumulate (Chaves *et al.,* 2009). The proline cycle requires NADPH to reduce glutamate to proline and, therefore, is supported by the electron transport chain. An enhanced proline accumulation in chloroplasts during stress conditions can maintain the NAPDH-NADP+ ratio and sustain the electronic flow rate between photosynthetic excitation centres (Hare and Cress, 1997). Furthermore, PDH genes, particularly P5CDH, are repressed under stress conditions to prevent proline degradation. Proline has been related to ROS scavenging activity since it is a singlet oxygen quencher (Matysik *et al.,* 2002). One of the effects of proline concerns the maintenance of photosystem II activity. Proline prevents the damaging effect of reactive oxygen species on the thylakoid membrane (Alia *et al.,* 1997). These effects were also highlighted in *Arabidopsis* exploiting mutants lacking the P5CS1 gene (Székely *et al.,* 2008). Our results are in accordance with previous literature and point out that the accumulation of proline is peculiar of stress conditions that include salinity; this may be explained by the need to limit photosystem II damage due to ROS species induced under stress conditions.

### Conclusions

This study investigated four single (salinity, heat, cold and drought) and combined stresses in plants at the phenotype, metabolomics, and stress markers levels. Except for salinity, each stress applied individually had a limited effect on growth and metabolism, while the cumulative impact of combined stresses was more detrimental to plants. This confirms that, by interacting, different stresses can have a synergic and stronger impact on plant health and performance, even in mild conditions and in the short term (Rillig *et al.,* 2019; Zandalinas *et al.,* 2021). This poses a new challenge for agricultural production in today’s climate change era. The distinctive interactions among stresses strongly corroborate the need to consider co-occurring stresses when the effects of climate change on agriculture are to be investigated.

Nonetheless, our findings pointed out a multi-layered effect of single vs. combined stresses, suggesting that using few markers may provide a limited picture of the processes underlying stress interactions. Consequently, more comprehensive approaches are advisable to unravel the additive effects among stresses, including non-targeted investigations. Another important outcome of this study is the confirmation that the stress level, timing of induction, and duration are pivotal conditions that must be accurately defined to investigate plant response to single or multiple abiotic stress. Besides paving the way towards future investigations, our results provide novel insights into the hierarchical effect of different stresses and their synergistic effects in plants.

## Abbreviations

DAS: days after sowing
CNTR: control
H: heat
D: drought
C: cold
S: salinity
DxH: drought x heat
DxC: drought x cold
SxH: salinity x heat
SxC: salinity x cold
ChlF: chlorophyll fluorescence kinetics
PAM: pulse amplitude modulated
MDA: malondialdehyde
HCA: hierarchical cluster analysis
OPLS-DA: orthogonal projection to latent structures discriminant analysis
VIP: variable importance in projection.

## Supplementary Material

***Supplementary Figure 1.*** OPLS-DA based on the metabolomic profile of *Arabidopsis thaliana* L. leaves (R^2^Y = 0.957; Q^2^Y = 0.735).

***Supplementary Figure 2.*** PlantCyc **(A)** metabolic pathway analysis and **(B)** details of secondary metabolism resulting from Volcano Plot analysis (FC>2, *P*<0.05).

***Supplementary Table S1*** RT, MS1 and MS/MS spectra of GABA, proline, betaine, and glycine, used for the targeted identification and quantification whit Quadrupole-Orbitrap MS.

***Supplementary Table S2.*** Morphological parameters (Leaf projected areas, Compactness and Roundness) of plants at T0-T4.

***Supplementary Table S3.*** Photosynthetic parameters (NQP, qP, Fm’/Fv’ and Fv/Fm) of plants at T0-T4.

***Supplementary Table S4.*** Putatively annotated compounds revealed with untargeted UHPLC-ESI7QTOF-MS analysis.

***Supplementary Table S5.*** Discriminant compounds (VIP score>1.2 and *P*<0.05) resulting from supervised OPLS-DA of all treatments taken together.

***Supplementary Table S6*** Compounds significant in at least one treatment (*P* <0.05, Benjamini correction) loaded for PlantCyc pathway analysis.

***Supplementary Table S7.*** Discriminant compounds (VIP score>1.2 and *P*<0.05) resulting from supervised OPLS-DA of D, DxH, DxC plants.

***Supplementary Table S8.*** Discriminant compounds (VIP score >1.2 and *P*<0.05) resulting from supervised OPLS-DA of S, SxH, SxC plants.

## Acknowledgements

This paper and related research have been conducted during and with the support of the Italian national inter-university PhD course in Sustainable Development and Climate change (www.phd-sdc.it) and of the PhD course in Agro-Food System, which founded the fellowships of ES and MADG, respectively. ACC was supported by Contrato-Puente from the Plan Propio of the University of Granada.

## Author contribution statement

LL: Conceptualization; ES, MADG: Formal Analysis; ES, MADG, ACC: Investigation; ES, MADG, ACC, LL: Methodology; ES, MADG: Visualization; ES, MADG, ACC, LL: Writing– Original Draft Preparation; LL: Writing – Review & Editing

